# Ecological niche contributes to the persistence of the Western x Glaucous-winged Gull hybrid zone

**DOI:** 10.1101/2023.12.14.571742

**Authors:** Xuewen Geng, Jeremy Summers, Nancy Chen

## Abstract

Hybrid zones occur in nature when populations with limited reproductive barriers overlap in space. Many hybrid zones persist over time, and different models have been proposed to explain how selection can maintain hybrid zone stability. More empirical studies are needed to elucidate the role of ecological adaptation in maintaining stable hybrid zones. Here, we investigated the role of exogenous factors in maintaining a hybrid zone between western gulls (*Larus occidentalis*) and glaucous-winged gulls (*L. glaucescens*). We used ecological niche models (ENMs) and niche similarity tests to quantify and examine the ecological niches of western gulls, glaucous-winged gulls, and their hybrids. We found evidence of niche divergence between all three groups. Our results best support the bounded superiority model, providing further evidence that exogenous selection favoring hybrids may be an important factor in maintaining this stable hybrid zone.

## Introduction

Hybridization is widely observed among related species, and the areas where lineages overlap (hybrid zones) offer a unique opportunity to investigate the role of ecology in speciation (Barton & Hewitt 1985; Abbott *et al*. 2013). In many cases, there is a balance between selection and gene flow within a hybrid zone, resulting in a stable hybrid zone over time (Moore 1977; Barton 1979). Both environmental variation and geographical barriers within a hybrid zone can facilitate the physical separation of the parental populations, potentially reinforcing existing levels of reproductive isolation and population divergence. At the same time, there are opportunities for hybrids with unique trait combinations or intermediate features to secure ecological niches different from their parental species along an environmental gradient (Endler 1977; Harrison 1993; Bugg 2007; Taylor *et al*. 2015).

Previous studies have proposed multiple models to explain hybrid zone stability. These models can be classified based on whether the selective forces acting on hybrids are endogenous or exogenous (Moore 1977). Endogenous selection refers to selection due to genetic incompatibilities independent of the environment, while exogenous selection refers to the differential selection of hybrids depending on the environment (Barton & Hewitt 1985). The tension zone model proposes that hybrid zones are maintained by an equilibrium between the dispersal of parental species into the hybrid zone and endogenous selection against hybrids (Key 1968; Moore 1977). The geographical selection-gradient model also states that hybrid zone stability is maintained by a balance between the dispersal of parental species and selection against hybrids, but here selection against hybrids is exogenous (Moore 1977; Barton & Hewitt 1985). The bounded superiority model argues that exogenous selection favors hybrids over either parental species because hybrids occupy a unique niche within the transitional zone (Moore 1977). Previous studies have found evidence supporting each of these three models in different hybrid zones (*e.g.*, Gay *et al*. 2008, Culumber *et al*. 2011 for the tension zone model; Edwards *et al*. 2015 for the geographical selection gradient model; Wang *et al*. 1997, Good *et al*. 2000, De La Torre *et al*. 2013 for the bounded superiority model). More studies of hybrid zone stability in other species are necessary to determine which of these models is more prevalent in nature.

One approach to identify the forces maintaining hybrid zone stability is to quantify the role of exogenous factors by comparing the niches of parental species and their hybrids (Swenson 2006; 2008). Ecological niche models (ENMs) use spatial environmental data and species occurrence localities to predict the possibility of species occurrence across different environments, providing estimates of the realized niche of the focal species (Guisan & Thuiller 2005; Broennimann *et al*. 2012). This approach has been used to investigate niche divergence and hybrid zone stability in multiple taxa, such as the hybrid zones between brown lemurs *Eulemur rufifrons* and *E. cinereiceps* (Johnson et al. 2016), tidal marsh birds *Ammodramus caudacutus* and *A. nelsoni* (Walsh et al. 2015), and swordtails *Xiphophorus birchmanni* and *X. malinche* (Culumber et al. 2012).

Here, we used ENMs to investigate the role of environmental variation in maintaining a well-studied hybrid zone between western gulls (*Larus occidentalis*) and glaucous-winged gulls (*L. glaucescens*) in the Pacific Northwest region of North America (Hoffman *et al*. 1978; Bell 1996; 1997). Although there is evidence that this hybrid zone has expanded since its discovery (Hoffman *et al*. 1978; Bell 1996; Good *et al*. 2000) and shifted south (Gay *et al*. 2008), cline analyses do not suggest the species are fusing (Gay *et al*. 2008). Thus, for the sake of our analysis in this study, we assume that the hybrid zone is stable or persistent over time (as in Megna *et al*. 2014). Previous studies of this hybrid zone have focused on comparing reproductive success, mating patterns, population structure, and clinal variation of the hybrids and the parental species (Hoffman *et al*. 1978; Larsen 1982; Speich & Wahl 1989; Bell 1992; Bell 1997; Good *et al*. 2000; Moncrieff *et al*. 2013; Megna *et al*. 2014). Results of these studies, however, support different models of hybrid zone stability (summarized in **Table 1**), thus which model best explains the stability of this hybrid zone remains up for debate. No previous studies of this hybrid zone have assessed the significance and contribution of potential differences between the ecological niches of the hybrids and the parental species to hybrid zone stability.

**Table 1:** Support for different models of hybrid zone stability from previous published work on the western x glaucous-winged gull hybrid zone and the present study.

| Study focus | Prediction | Model | Outcome |
| --- | --- | --- | --- |
| Hybrid reproductive success | 1) Hybrids exhibit lower reproductive success compared to parental species | Tension zone model or geographical selection-gradient model | **No.** Reproductive success of hybrids is equal to or higher than that of the parental species. |
|  | 2) Hybrids perform equal or better than at least one of the parental species | Bounded superiority model | **Yes.** Hoffman *et al.* 1978, Good 1998 and Good *et al.* 2000 suggest hybrids perform better than the parental species. Moncrieff *et al.* 2013 and Megna *et al.* 2014 suggest hybrids and the parental species perform equally. |
| Assortative Mating | 1) Assortative mating exists because hybrids are being selected against: Mating with hybrids results in a decrease in reproductive success. | Tension zone model or geographical selection-gradient model | **Maybe.** Hoffman *et al.* 1978, Bell 1997, and Megna *et al.* 2014 all find evidence of assortative mating. However, Megna *et al.* 2014 find there is no decrease in reproductive success for dissimilar pairs, and Good *et al.* 2000 also find no significant correlations between the hybrid indices of male and female gulls. |
|  | 2) Preference for hybrid mates | Bounded superiority model | **No.** No evidence has been found showing a preference for mating with hybrids. |
| Population density in the hybrid zone | 1) Low density of hybrids within the hybrid zone | Tension zone model or geographical selection-gradient model | **No.** Hybrids are prevalent within the hybrid zone. |
|  | 2) Higher density of hybrids in the hybrid zone compared to outside the hybrid zone | Bounded superiority model | **Yes.** Hoffman *et al.* 1978 and Bell 1992 suggest that hybrids occur more frequently within the hybrid zone. |
| Cline shape | 1) Clines are sigmoid and narrow in shape and coincident | Tension zone model or geographical selection-gradient model | **Yes.** Gay *et al.* 2008 find sigmoid and narrow clines in phenotypic traits and molecular markers. |
|  | 2) Clines have more variable shape and noncoincident | Bounded superiority model | **Maybe.** Bell 1996 found shallow clines at allozyme markers |
| Ecological niche | 1) Niche conservatism between all three groups | Tension zone model | **Maybe.** Range-breaking tests find no significant geological boundary between all three groups, but these results could simply indicate a mosaic hybrid zone (this study). |
|  | 2) Niche conservatism between the parental species and the hybrids, and niche divergence | Geographical selection-gradient model | **No.** There is no evidence of niche conservatism between the parental species and hybrids but niche divergence between the |
|  | between the parental species |  | parental species. |
|  | 3) Niche divergence between all three groups | Bounded superiority model | **Yes.** Niche identity tests and response curves from ENMs indicate niche divergence among all three groups (this study). |

In this study, we tested whether hybrid gulls exhibit a different niche than their parental species and how environmental variation may contribute to the distribution and stability of the hybrid zone. We constructed ENMs, characterized environmental variation associated with the distributions of glaucous-winged gulls, western gulls, and hybrid gulls, and quantified the differences between the niches occupied by the two parental species and their hybrids. We hypothesized that if there is no niche divergence between the parental species and the hybrids or between the two parental species and the occurrence of hybrids is highly associated with the presence of parental species, then exogenous factors do not contribute to the stability of this hybrid zone, indicating support for the tension zone model. If the hybrid zone is best explained by the geographical selection-gradient model, we hypothesized that we should observe niche divergence between the two parental species, but not necessarily between the hybrids and the parental species, as selection is assumed to be against the hybrids. Instead, if the hybrid zone conforms to the bounded superiority model, we predicted that there should be niche divergence between all three groups.

## Materials and Methods

### Study System

We studied the western x glaucous-winged gull hybrid complex along the western coastline of North America. Western gulls breed from central Baja California north to Washington, and glaucous-winged gulls breed further north, from Washington to Alaska (**Figure 1**; Birds of the World 2023). The hybrid zone is considered to be a narrow ecotone along the Washington coastline (Reagan 1911; Hoffman et al. 1978; Birds of the World 2023). Hybrids between western gulls and glaucous-winged gulls were first noted in the early 20th century, and are prevalent within the hybrid zone, occurring in higher frequencies than the parental species at some locations (**Figure 1**; Dawson 1908; Dawson *et al*. 1909; Bell 1992). The hybrids and the parental species can be distinguished primarily based on their mantle and wingtip plumage color: western gulls have darker grey plumage, glaucous-winged gulls have lighter grey plumage, and hybrids have an intermediate shade of grey. Other distinguishing traits include iris color, orbital ring color, and beak color (Bell 1996; Bell 1997; Moncrieff 2013). These three taxonomic groups can be visually distinguished in the field, allowing us to use citizen science databases for data acquisition (see below).

**Figure 1:**
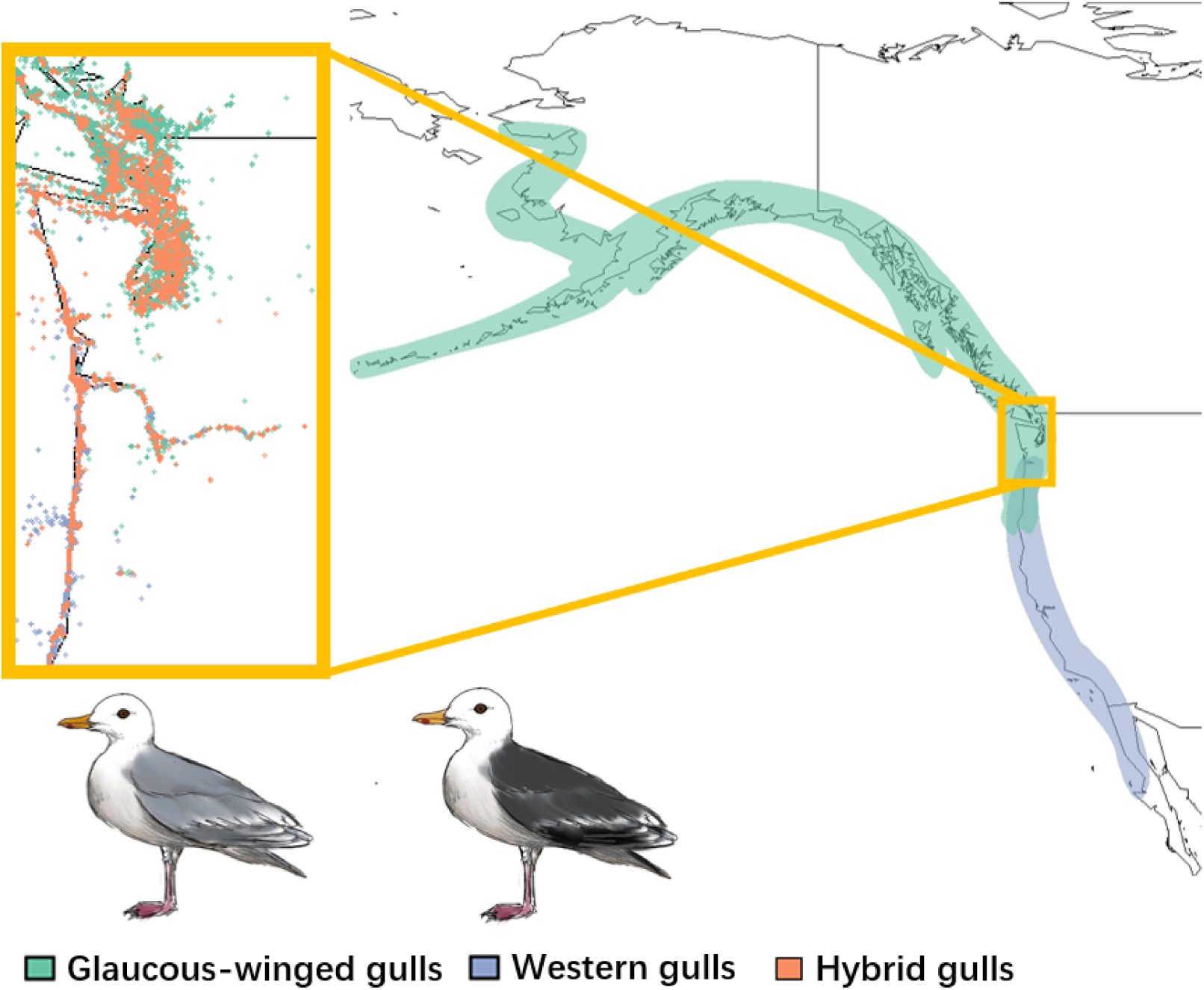
The location of the hybrid zone and the ranges of glaucous-winged gulls (*Larus glaucescens*; green), western gulls (*Larus occidentalis;* blue), and their hybrids (orange) in North America. The inset shows eBird occurrence data for the two parental species and hybrid individuals in the hybrid zone. Western gulls and glaucous-winged gulls can be visually distinguished best by differences in mantle and wingtip plumage color.

### Species Occurrence Data

ENMs use both species occurrence data and environmental layers to predict the niche of a species. We obtained occurrence data for the parental species and the hybrids from the citizen science database eBird (Sullivan *et al*. 2009). eBird is the world’s largest citizen science database for bird occurrence records, allowing birders from all around the world to document the distribution, abundance, and identities of the birds they encounter. eBird has a robust review process for ensuring species identity. To enter a record as a hybrid, eBird requires individuals to submit supporting materials such as photos, audio, field notes, or video evidence, which is then verified by local eBird reviewers (Sullivan *et al*. 2009). Here, we used the eBird Basic Dataset version EBD-relMay-2023.

We used the *auk* package in R to download and filter eBird data (Strimas-Mackey *et al*. 2022). Since western gulls and glaucous-winged gulls are known to be partially migratory, we extracted eBird occurrence data within the breeding season (defined as June to July based on the temporal overlap of their breeding seasons; Murphy *et al*. 1984; Vermeer *et al*. 1988; Annett *et al*. 1989; David *et al*. 2015). In addition, as hybrid gulls were not frequently reported until 2010, we only included occurrence points recorded between 2010-2023. To lessen sampling effort bias, we filtered observations based on survey protocols (*Stationary* and *Traveling* checklists only), duration (< 360 minutes), distance traveled (< 10km), time of the day (6:00 – 21:00), and the number of observers (<=10 people) for each observation based on *Best Practices for Using ebird Data* (Strimas-Mackey *et al*. 2023). We also only included checklists marked as complete (ones for which birders indicate they recorded every bird they detected) to reduce the impact of taxonomic preferences and bias in detection (Johnston *et al*. 2019). We set the spatial extent based on the longitudes and latitudes of the occurrence points:114° W-175° W, 31° N-62° N (Hoffman *et al*. 1978; Barton & Hewitt 1985; Bell 1996; 1997; **Figure 1**). We then manually removed outliers. Our filtered eBird dataset included 81,363 observations of western gulls, 66,697 observations of glaucous-winged gulls, and 9,419 observations of hybrid individuals. The occurrence data span seven states across three countries (Canada, the United States, and Mexico), including Baja California (Mexico), British Columbia (Canada), Yukon Territory (Canada), California (United States), Oregon (United States), Washington (United States), and Alaska (United States), which fits with current understanding of species ranges for these gulls.

To account for potential sampling effort bias introduced by geographic factors, we performed spatial thinning using the *spThin* package in R (Aiello-Lammens 2015). We chose a thinning grid of 0.5 km x 0.5 km because previous studies showed this grid size efficiently removes redundant records while including spatially valuable data (Steen *et al*. 2020). We then balanced the sample sizes for each species and the hybrids by randomly sampling records from the thinned dataset. Our final dataset consisted of 1,500 records for each taxonomic group (**Figure S1**).

### Environmental Data

We included a total of eight environmental variables in our ENMs. We obtained bioclimactic variables from WorldClim (Fick & Hijmans 2017), landcover data from the North American Land Change Monitoring System (NALCMS; Homer *et al*. 2017), elevation data from *elevatr* package in R (Hollister *et al*. 2021), and distance from coastline data NOAA National Ocean Service (Stumpf & Kuring 2009). WorldClim is composed of a set of gridded climate layers with variables related to temperature and precipitation (Fick & Hijmans 2017). We downloaded the 19 bioclimatic layers from WorldClim using a 2.5-minute spatial resolution. We only included annual measures and excluded the isothermality layer, which is inherently highly correlated with other layers since it is calculated as the mean diurnal range layer divided by the temperature annual range layer. We downloaded the land cover type layer from NALCMS using a 30-m spatial resolution. This layer includes 19 landcover types that are jointly identified by government agencies from Canada, US, and Mexico. We labeled spaces along the shoreline that are covered by bioclimatic layers but missing landcover data as landcover type 18, water. We downloaded the world elevation map from the *elevatr* package in R. This package provides a combination of world elevation maps based on publicly accessible remote sensing maps from different countries. We downloaded the distance from the coastline map from NOAA National Ocean Service, which provides a world map of distance from the coastline with an uncertainty of 1 km. Locality grids marked with values <0 represent localities corresponding to the land, and localities marked with values >0 represent localities corresponding to the ocean.

We then extracted environmental data for each occurrence point from the thinned dataset and tested the correlation between different layers using the *vifstep* and *vifcor* tools from the *usdm* package in R (Naimi *et al*. 2014). These are two different ways of detecting collinearity using a variance inflation factor (VIF). *Vifstep* lists variables that yield a higher VIF than the threshold (10), whereas *vifcor* removes the variable that yields a higher VIF from a pair of variables that has a greater linear correlation than a specified threshold (we used 0.9; Naimi *et al*. 2014). Using both methods, we removed environmental layers that are highly correlated with other layers. Our final environmental variables were annual mean temperature, mean diurnal range, temperature seasonality, annual precipitation, precipitation seasonality, landcover type, elevation, and distance to the coastline.

### Environmental niche modeling

We used the maximum entropy method in Maxent to construct ENMs. Maxent predicts the suitability of environmental conditions for the species of interest based on species occurrence localities, background points, and environmental layers (Philips & Dudík 2008). The final suitability map produced from Maxent represents the probability of occurrence that contains the maximum entropy, or the most spread out distribution (Elith *et al*. 2010). Maxent offers a few advantages that are important to our study: (1) It uses presence-only data, avoiding potential biases introduced by predicted absence data from complete ebird records (Johnston *et al*. 2021). (2) It can consider both continuous and categorical environmental variables. (3) Maxent outputs a continuous probability of suitability raster that allows further comparative analyses between populations (Philips *et al*. 2006). The one disadvantage, however, is Maxent models are sensitive to sampling bias introduced by potential correlations between sampling efforts and specific environmental variables (Merow *et al*. 2013; Johnston *et al*. 2021). To correct for potential sampling effort bias introduced by spatial features across our spatial extent, we used the target-group background method to select background points. The target-group background method uses the occurrence points of target-related species sampled by the same methods within the zone of interest as background datasets to account for sampling bias (Ponder *et al*. 2001; Philips *et al*. 2009; Vollering 2019). This approach has been proven to be effective in Maxent models (Barber 2022). Thus, we downloaded, filtered, and thinned the eBird Basic Dataset (version EBD-relMay-2023) for all species using the same protocol we used for the parental species and hybrids. We then extracted 10,000 points from the filtered all-species dataset within our study extent to serve as our target-group background points.

We used the *maxent* package in R with our eight environmental variables, species occurrence localities, and target-group background points as input (Jurka 2012). We set landcover type as a categorical variable and the other seven variables as continuous variables. In addition, we cross-validated each model 10 times based on a 20% testing and 80% training percentage to calculate confidence intervals. To evaluate model performance, we calculated the area under the receiver operating curve (AUC) for each model. An AUC value of 0.5 indicates the model performs similar to random prediction, and an AUC value of 1 means the model has perfect prediction power. Typically, an AUC value above 0.7 indicates reliable performance (Metz 1978; Swets 1988).

We analyzed the importance of each environmental variable by evaluating their percent contribution and permutation importance. Both are measures of variable contribution and are automatically produced by the *maxent* package. The *maxent()* function calculates the percent contribution based on the contribution of variables for the final optimal model. The program calculates the permutation importance of each variable based on changes in AUC values when permuting a specific variable. We calculated the mean and confidence intervals using results over 10 iterations (Philips & Dudík 2008, Elith *et al*. 2011, Philips *et al*. 2017).

To test the accuracy of our eBird models, we tested model transferability between eBird models and maxent models built on another citizen science database: The North American Breeding Bird Survey (BBS). Please see Appendix 1 for more.

### Niche similarity tests

To test for differences in ecological niches between each of the parental species and the hybrids, we performed niche identity tests and pairwise blob range-breaking tests using the *ENMTools* package in R (Warren *et al*. 2021). The niche identity test looks for a significant difference between two ecological niche indices (Schoener’s *D* and Warren’s *I*) calculated for randomized controls and the population of interest (Schoener 1968; Warren *et al*. 2008). The range-breaking test uses the same set of indices to check for a distinct boundary between the occurrence points of two populations (Warren *et al*. 2008). Schoener’s *D* calculates the overlap in niches between two populations by summing the absolute differences in the relative use of a particular type of habitat:

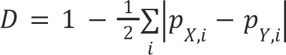

where *p_X_* and *p_Y_* represent the probability of occurrence for species *X* and *Y* respectively, and *i* represents each grid of the study extent. Warren further revised this equation by integrating Hellinger’s distance *H* to develop Warren’s *I* (Warren *et al*. 2008):

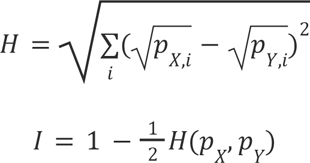

Both indices are commonly used in quantifying niche overlap. The values of both indices range from zero, indicating no niche overlap, to one, indicating a perfect niche overlap between each pair of populations tested. We used a similar approach for both the niche identity test and the range-breaking test: We tested our original population of interest and 100 iterations of the randomized controls, which we randomly chose from the original population. Then we compared the niche indices of our original population of interest and the control populations to calculate significance. The only difference between these two tests is that the range-breaking test extracts the control population by first randomly selecting one point and then adding points surrounding this origin (Warren *et al*. 2021). We used a significance threshold of 0.05.

## Results

To quantify the ecological niches of parental and hybrid gulls, we built ENMs for glaucous-winged gulls, western gulls, and their hybrids (**Figure 2**). The mean AUC of our models are all above 0.7 (glaucous-winged gulls: mean=0.891, SD=0.009; western gulls: mean=0.906, SD=0.006; hybrids: mean=0.898, SD=0.010), indicating our models can reliably predict occurrence (Hosmer & Lemeshow 2000). The predicted species distributions from our ENMs match the known distributions: glaucous-winged gulls prefer northern habitats and western gulls prefer southern habitats, with an overlap in the middle where the hybrids appear (**Figure 1**). The hybrid distribution extends beyond the historical recorded hybrid range, but with the highest probabilities of occurrence along the Washington coastline, where most previous studies have found and studied them.

**Figure 2:**
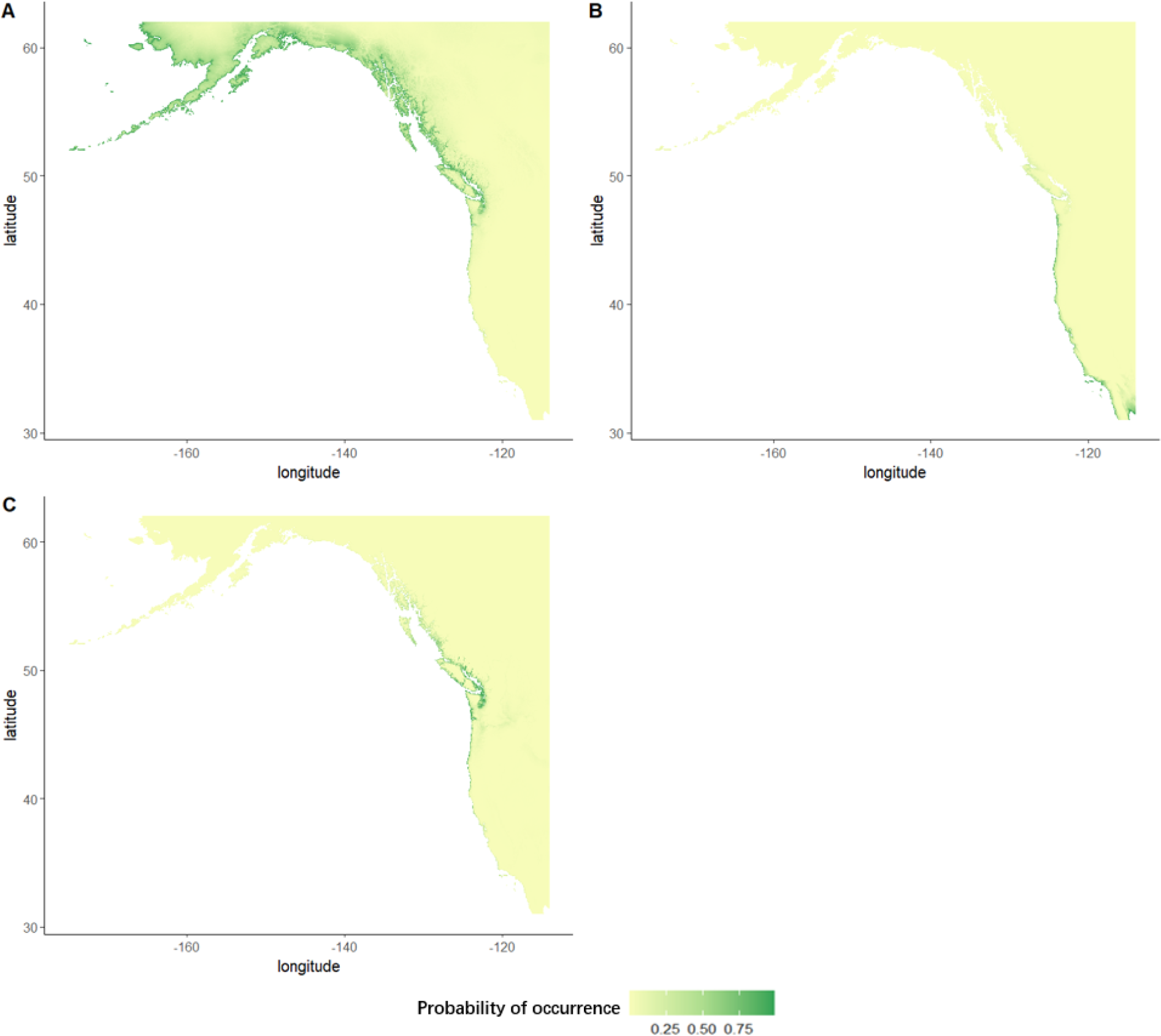
The predicted probability of species occurrence from maxent ENMs for (B) glaucous-winged gulls, (B) western gulls, and (C) their hybrids. Green represents a higher possibility of occurrence and yellow represents a lower possibility of occurrence

We calculated the niche indices Schoener’s *D* and Warren’s *I* (**Table 2**) and performed a niche identity test to test for significant niche divergence between the parental species and the hybrids. We found that glaucous-winged gulls and hybrid gulls have significantly different niches (*p* < 0.01; **Figure 3A**), with the randomized controls yielding indices that are significantly higher than the testing populations. We obtained similar results between western gulls and the hybrids, and between the two parental species (*p* < 0.01 for both tests; **Figure 3B-C**). These results suggest that all three populations have different niches relative to the null expectations.

**Figure 3:**
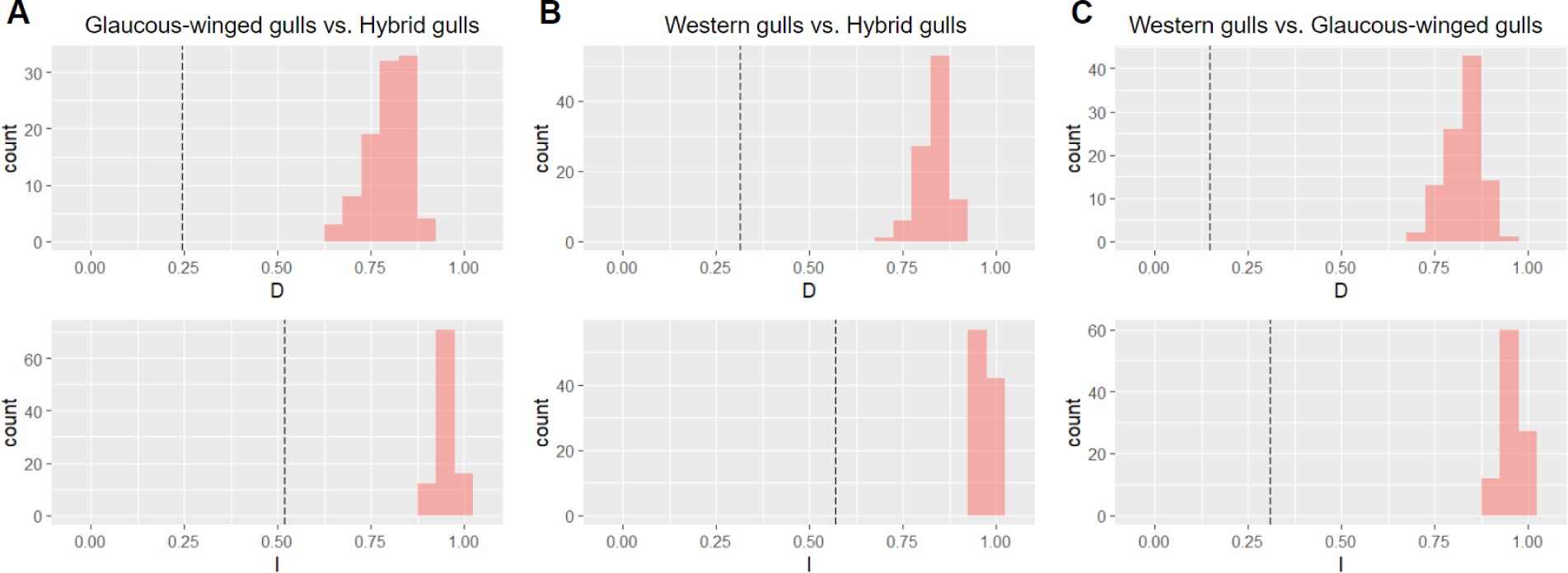
Niche identity test results between (A) glaucous-winged gulls and hybrid gulls, (B) western gulls and hybrid gulls, and (C) glaucous-winged gulls and western gulls. Results for Schoener’s *D* are on the top row and Warren’s *I* on the bottom. Histograms show the null distribution, and the dotted black line is the observed value. All tests show significantly lower-than-expected niche indices, indicating significant niche divergence between all pairwise comparisons.

**Table 2:** Niche indices calculated for the niche identity test and range-breaking test between western gulls, glaucous-winged gulls and their hybrids.

|  | **Glaucous-winged gulls vs. Hybrid gulls** | **Western gulls vs. Hybrid gulls** | **Glaucous-winged gulls vs. Western gulls** |
| --- | --- | --- | --- |
| **Schoener's *D*** | 0.248 | 0.315 | 0.146 |
| **Warren's *I*** | 0.520 | 0.570 | 0.309 |

We performed range-breaking tests to examine whether there is a distinct boundary between the parents and the hybrids within the hybrid zone and whether physical geological barriers contribute to niche divergence. We found that the empirical indices are not significantly lower than the randomized controls for glaucous-winged gulls and the hybrids in the range-breaking test (*p*<0.01; **Figure 4A**), suggesting that there is no distinct boundary between these two populations. We obtained similar results between western gulls and the hybrids and between the two parental species (*p*<0.01 for both tests; **Figure 4B-C**). These results suggest that there is no abrupt geographical boundary associated with sudden environmental gradients in this hybrid zone.

**Figure 4:**
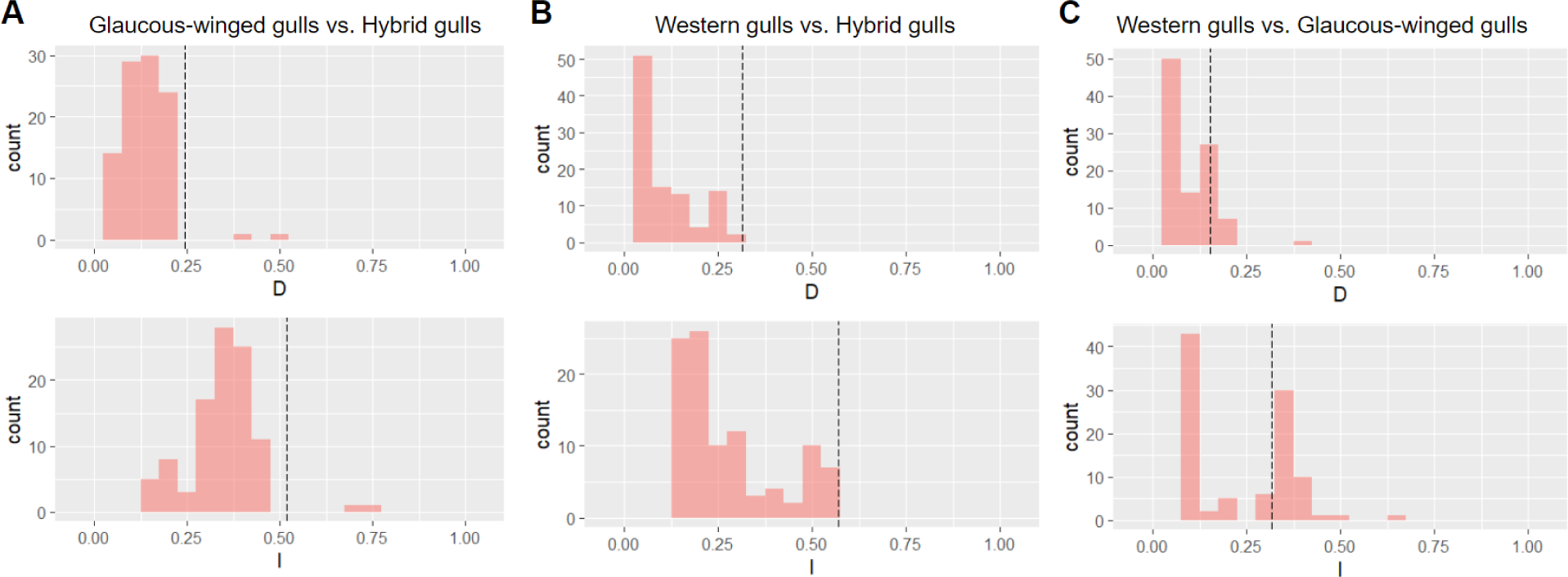
Range-breaking test results between (A) glaucous-winged gulls and hybrid gulls, (B) western gulls and hybrid gulls, and (C) glaucous-winged gulls and western gulls. Results for Schoener’s *D* are on the top row and Warren’s *I* on the bottom. Histograms show the null distribution, and the dotted black line is the observed value. The observed value is not significantly different from the null expectation for any test. Thus there is no evidence for any significant range boundaries between populations.

We examined how environmental variation is associated with species occurrence data by analyzing the response curves and individual variable contributions from our ENMs. We found that different variables contributed the most to the probability of occurrence across the three taxa: isothermality (64.6% contribution relative to the maxent model; 40% permutation importance) and mean diurnal range (24.4%; 51%) contributed the most for western gulls, mean diurnal range (59.9%; 46.3%) and landcover types (24%; 28.1%) for glaucous-winged gulls, and mean diurnal range (41.5%; 52.6%) and annual mean temperature (22.9%; 18.5%) for the hybrids (**Figure 5**). These results further suggest that western gulls, glaucous-winged gulls, and hybrids occupy different ecological niches.

**Figure 5:**
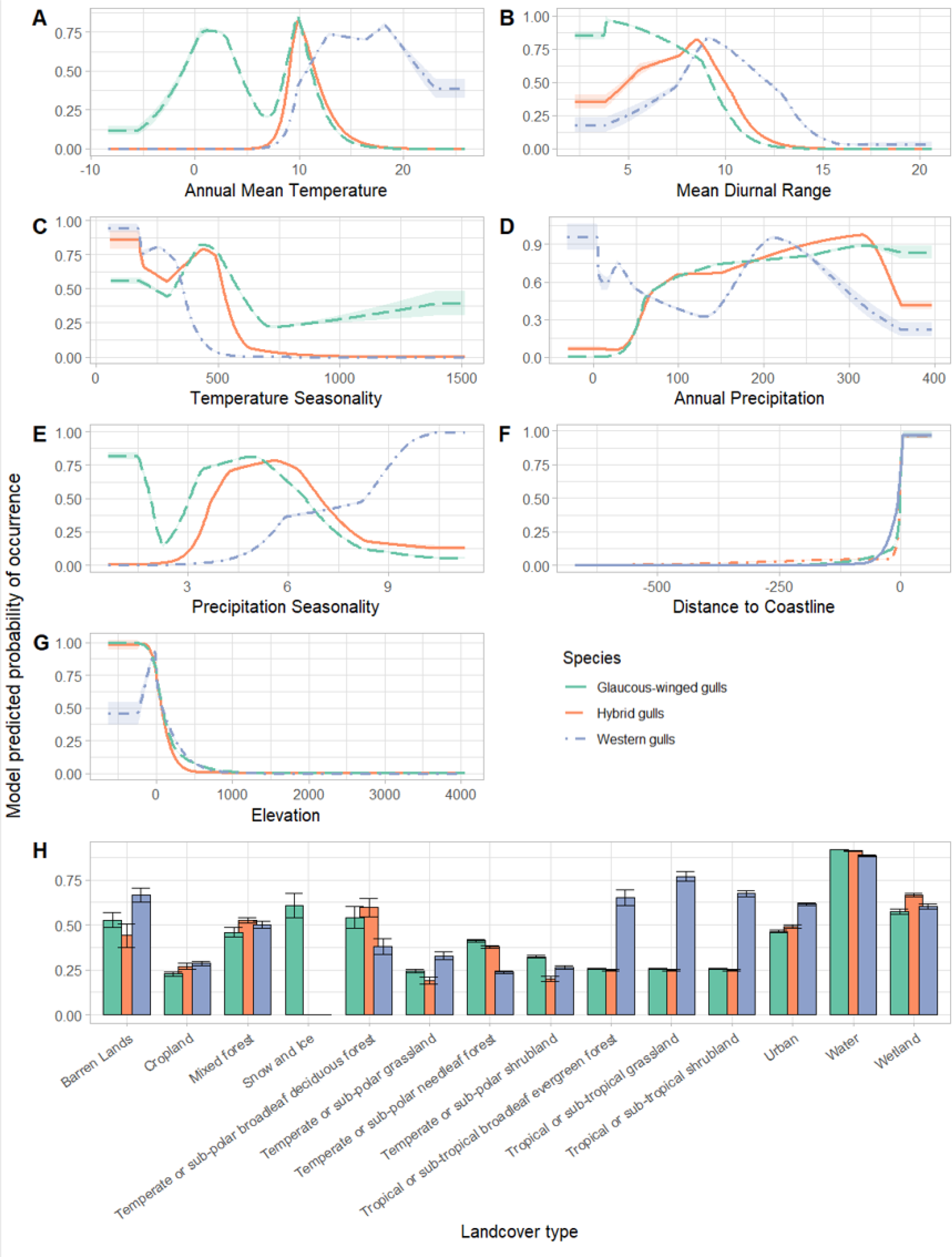
Response curves in relation to (A) annual mean temperature, (B) mean diurnal range, (C) temperature seasonality, (D) annual precipitation, (E) precipitation seasonality, (F) distance to coastline, (G) elevation, and (H) landcover type for glaucous-winged gulls (green), western gulls (blue), and hybrid gulls (orange). Confidence intervals were calculated across 10 iterations of the models.

## Discussion

Our results indicate that environmental gradients within the hybrid zone between western gulls and glaucous-winged gulls may contribute to maintaining hybrid zone stability under the bounded superiority model. The bounded superiority model predicts both niche divergence between the two parental species and niche divergence between the parental species and the hybrids. Our niche identity tests show that western gulls, glaucous-winged gulls, and their hybrids all occupy significantly different ecological niches. The differences in the response curves and contributions of individual environmental variables suggest that differences in isothermality, landcover type, mean diurnal range, and annual mean temperature may underlie the observed niche divergence. In addition, differences in variable contributions and response curves among the three populations suggest there are differences in habitat preferences.

Under the tension zone model, there should be niche conservation within the hybrid zone among all three gull populations. Our range-breaking tests found no distinct boundaries between populations. This result disagrees with our niche identity test results and could be explained by the complex habitat preferences of gulls (Hoffman *et al*. 1987; Bell 1992). These three populations show differences in the importance of environmental variables and response curves based on their ENMs. Niche diversification between the hybrids and the parental species could be caused by variation in habitat preference along a smoother but more mosaic environmental gradient instead of a steep environmental gradient (**Figure 5**), which would result in a lack of distinct boundaries between parental species and hybrid distributions detectable by a range-breaking test. Thus the range-breaking test results do not necessarily indicate niche overlap among the three populations. The geographical selection-gradient model expects niche divergence between the parental species but not between the hybrids and the parental species, (Moore 1977). We found no support for this model from either the range-breaking test or the niche identity test: the range-breaking test suggests a lack of distinct boundaries among the three populations whereas the niche identity test suggests niche divergence among the three populations.

Previous studies of this hybrid zone have investigated mating success, hatching success, or fledging success to compare the fitness of the hybrids with that of the parental species (**Table 1**). However, these studies found contradicting results: some studies showed that hybrids outperform the parental species significantly within the hybrid zone (Hoffman *et al*. 1978, Good 1998; Good *et al*. 2000), supporting the bounded superiority model, but others found that the reproductive performance of the hybrids is not significantly different from that of at least one of the parental species (Bell 1996, Moncrieff *et al*. 2013, Megna *et al*. 2014), finding no strong support for any of the three models. Assortative mating based on appearance has been observed within multiple sites in both the parental species and the hybrids (Moncrieff *et al*. 2012; Moncrieff *et al*. 2013; Megna *et al*. 2014). However, as hybrid gulls display appearances that span a wide spectrum between the two parental species, it can be difficult to interpret the causes and results of this behavior (Hoffmann *et al*. 1978; Good *et al*. 2000; Bell 1996; 1997; Moncrieff *et al*. 2013; Megna *et al*. 2014). Phenotypic and genotypic cline analyses found both steep, stepped, and more random shape cline models for different features of the gulls, and provided support for either the tension zone model or the bounded superiority model (Bell 1996; Gay *et al*. 2008). Our study adds to previous work by using ecological niche modeling to assess niche differences between taxa in this hybrid zone, and our results provide additional support for the bounded superiority model.

We tried to address or account for multiple concerns about predicting ecological niches in this study. (1) The presence-background nature of maxent models may introduce potential inaccuracy in predicting unsuitable habitats, as absence data is not considered in the model (*e.g.*, Svenning & Skov 2004, Jiménez-Valverde *et al*. 2008). However, previous work suggests that part of the information contained in absence data is also available in presence data, and the problem of false absences in presence-absence modeling can be more detrimental to niche prediction than using presence-only data (Elith *et al*. 2011). (2) We acknowledge that identifying gull species in the field can be challenging, and it is possible that the eBird occurrence data included identification errors. Although the stricter policy of inputting hybrid records eliminates some identification errors, we cannot fully eliminate cases in which a hybrid is misidentified as one of the parental species. We used species occurrence data from another data source, the North American Breeding Bird Survey (BBS), and performed a model transferability test to check if ENMs are consistent across different sources (**Appendix 1**). The resulting high AUC values (> 0.75; **Appendix 1**) provide validation for the accuracy of our maxent models. (3) Our analyses consider the comparatively larger ranges of the parental species outside of the hybrid zone, which could have influenced the accuracy of our models in predicting the ecological niches of the parental individuals within the current known hybrid zone (Lee-Yaw *et al*. 2016). However, as gulls can travel long distances within days, only considering individuals within or near the hybrid zone will also influence the accuracy of niche estimation. Including the whole range of both parental species can help us to better understand the overall niche of each species and the estimated potential overlap between the range for hybrids and parental species. (4) Potential bias in weighting variables may be introduced if the environmental variables are correlated with each other (Journé *et al*. 2019). We tried to eliminate the influence of such bias by removing highly correlated environmental variables, only keeping 8 of 22 variables. (5) Finally, one limitation of our work is we only included abiotic environmental variables in our models. Future studies are needed to characterize the potential influence of biotic factors on the stability of this hybrid zone.

## Conclusion

Our study provides additional insights into the role of environmental variation in maintaining the western x glaucous-winged gull hybrid zone. Using ENMs fitted with data from citizen science databases, we characterized the ecological niches of the parental species and the hybrids and showed that all three populations occupy different niches. Our results best support the bounded superiority model, suggesting that hybrid gulls are better adapted to the environment in the hybrid zone than either of the two parental species, which aligns with previous studies that observed higher reproductive success of hybrid individuals. Additional empirical support for the bounded superiority model indicates the potential importance of environmental variation in the maintenance of hybrid zone stability.

## Data Accessibility Statement

All metadata and code used in this study are available on GitHub: https://github.com/ggg80/HybridGull_Repo.

## Author Contributions Section

**Xuewen Geng:** Conceptualization (lead); data curation (lead); formal analysis (lead); methodology (lead); visualization (lead); writing – original draft (lead); writing – review and editing (equal). **Jeremy Summers:** Conceptualization (equal); methodology (equal); supervision (equal); visualization (support); writing – review and editing (equal). **Nancy Chen:** Conceptualization (equal); funding acquisition (lead); methodology (equal); project administration (lead); supervision (lead); visualization (support); writing – review and editing (equal).

## Acknowledgments

We thank the other members of the Chen lab for providing helpful feedback. JS was supported by NIH grant 1R35GM133412 to NC.

