## Supplemental Materials for "Ecological niche contributes to the persistence of the Western x Glaucous-winged Gull hybrid zone"

### Supplementary Figures

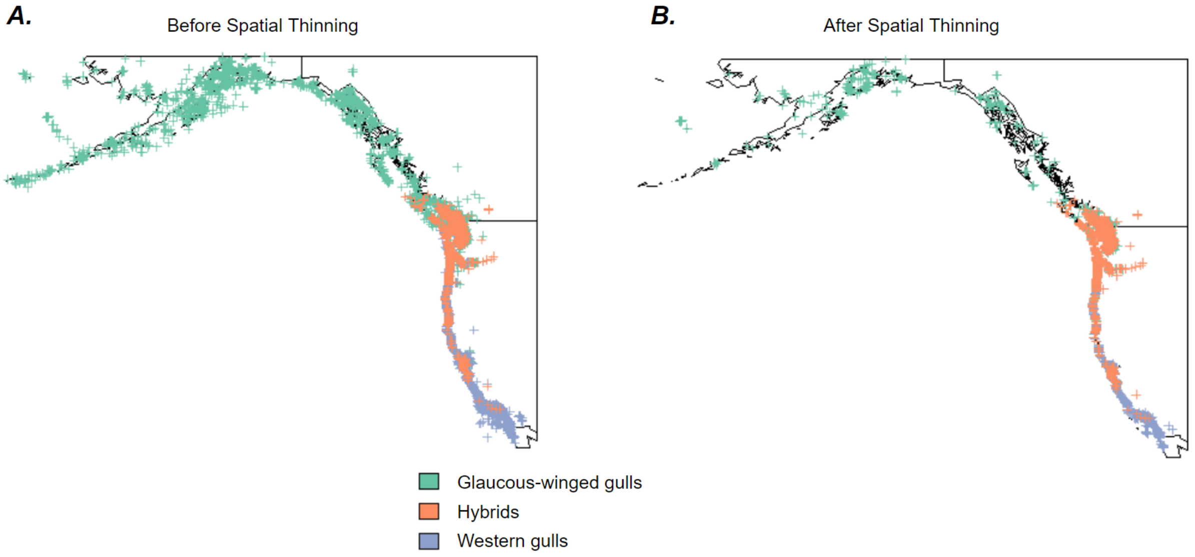


**Figure S1:** Species occurrence points for glaucous-winged gulls (green), western gulls (blue), and hybrid gulls (orange) (A) before and (B) after spatial thinning.

**Appendix 1. Model Transferability Test**

As gulls are notorious for their physical similarities across species and challenging species identification, we also constructed ENMs using another citizen science database, the American Breeding Bird Survey (BBS), to validate our eBird models using the model transferability test (Wioletta *et al.* 2016; Ziolkowski *et al.* 2022). BBS is a long-term breeding bird population and distribution monitoring program that records species occurrence and abundance along more than 4000 routes each year by bird experts (Ziolkowski *et al.* 2022). We extracted occurrence data from the same study extent using the same filters and constructed Maxent models using the same settings as the eBird data (see Materials and Methods). We did not have enough *L. occidentalis-glaucescens* hybrid occurrence records in the BBS dataset to fit ENMs (there were only two occurrence points), so we only considered the two parental species (*L. occidentalis* and *L. glaucescens*).

To validate our ENMs for accuracy, we used model transferability tests to determine whether our ENMs trained on eBird data can predict distributions based on BBS occurrence data. Model transferability refers to the ability of a model to predict reliable results in unsampled geographical or temporal regions, or the consistency of the results across models built on different datasets (Peterson 2007; Wioletta *et al.* 2016). We used the *evaluate()* function from the *maxent* package in R to calculate the AUC value between models built on one of the two databases with testing data from the other database (Jurka 2012).

The resulting AUC values between each pair of parental species models are high (0.904 for glaucous-winged gulls; 0.915 for western gulls), which suggests that our models are consistent between different databases. We considered this result a validation of the accuracy of species identification of the eBird data we used in this study.

**Appendix 2. Code**

Please refer to this GitHub repository for all code used in this manuscript:
**https://github.com/ggg80/HybridGull_Repo**
